# Decellularised Human Liver is too Heterogeneous for Designing a Generic ECM-mimic Hepatic Scaffold

**DOI:** 10.1101/073270

**Authors:** Giorgio Mattei, Chiara Magliaro, Andrea Pirone, Arti Ahluwalia

## Abstract

Decellularised human livers are considered the perfect ECM (extracellular matrix) surrogate because both 3-dimensional architecture and biological features of the hepatic microenvironment are thought to be preserved. However, donor human livers are in chronically short supply, both for transplantation or as decellularised scaffolds, and will become even scarcer as life expectancy increases. It is hence of interest to determine the structural and biochemical properties of human hepatic ECM to derive design criteria for engineering bio-mimetic scaffolds. The intention of this work was to obtain quantitative design specifications for fabricating scaffolds for hepatic tissue engineering using human livers as a template. To this end, hepatic samples from 5 human donors were decellularised using a protocol shown to reproducibly conserve matrix composition and micro-structure in porcine livers. The decellularisation outcome was evaluated through histological and quantitative image analyses to evaluate cell removal, protein and glycosaminoglycan content per unit area. Applying the same decellularisation protocol to human liver samples obtained from 5 different donors yielded 5 different outcomes. Only 1 liver out of 5 was completely decellularised, while the other 4 showed different levels of remaining cells. Moreover, protein and glycosaminoglycan content per unit area after decellularisation were also found to be donor-dependent. The donor-to-donor variability of human livers thus precludes their use as templates for engineering a generic 'one-size fits all' ECM-mimic hepatic scaffold.

## 1 Introduction

The liver is the largest visceral organ in the human body. Although it has a high regenerative potential, drugs, toxins or viral infections can cause extensive damage to hepatocytes, reducing tissue function and compromising regeneration. The World Health Organization (WHO) report shows that 2.5% of total deaths in the world are because of liver disease (Lopez et al. 2006), and this will become the 14^th^ most common cause of death by 2030 (Mathers and Loncar 2006).

Extensive damage to hepatocytes can lead to liver failure, which is likely to cause subsequent comorbidities, leading to neurological impairment, renal dysfunction, haematological disturbance and metabolic abnormalities. Usually death occurs as a result of multi-organ failure and, to date, allogeneic liver transplantation is the treatment of choice for patients with end-stage liver disease. However, it is limited by both the high cost and the severe shortage of donor organs (Schramm et al. 2010), thus prompting the demand of new alternative strategies to treat liver-diseased patients. Living donor organ transplantation, employing partial grafts rather than whole cadaveric livers, can help cope with the shortage of donor organs, but it is still not able to overcome the requirement for donor livers (Adam et al. 2012). Alternative strategies, such as whole-organ xenotransplantation (Ekser et al. 2009), hepatocyte transplantation (Strom et al. 1997), extracorporeal devices (Rifai et al. 2003) and stem cell therapies (Ishii, Yasuchika, and Ikai 2013), are being pursued throughout the world either to prolong the life of a patient until a donor liver is available or to regenerate functional hepatic tissue, but have had limited clinical success (Struecker, Raschzok, and Sauer 2014; Wertheim, Baptista, and Soto-Gutierrez 2012). Methods to obtain bio-artificial tissue via decellularisation of healthy human liver unsuitable for transplant have been proposed. For instance, Mazza et al. recently decellularised whole human liver through vascular perfusion, reporting minimal adverse effects on mice subcutaneously implanted with small pieces of human derived ECM (Mazza et al. 2015). However, despite the improvement of donor screening and transplantation procedures, which resulted in 87.4% of donor livers being successfully transplanted last year in Italy (“https://trapianti.sanita.it/statistiche/PEorg.asp” 2015), healthy human livers suitable for whole organ or split transplant, let alone for whole organ decellularisation, continue to be in short supply. A change in this trend is highly unlikely, given the demographic shift and decreases in mortality rates in developed countries as well as the development of new techniques for prolonging organ preservation after death. Therefore, methods that do not rely on the use of healthy human tissue (be it for transplantation or as decellularised scaffolds) are needed to overcome the shortage of donor organs.

Tissue engineering and regenerative medicine approaches are widely investigated to develop functional liver constructs for both in-vitro and in-vivo applications (Bhatia et al. 2014; Ananthanarayanan et al. 2011). These methods are based on the successful interaction between three components: i) cells of the target tissue, ii) a scaffold serving as a supportive template for cells promoting their adhesion, proliferation and development, and iii) a bioreactor for the dynamic cultivation of the cell-scaffold construct providing cells with appropriate physicochemical cues to direct them towards the expression of the desired tissue phenotype (Pörtner and Giese 2006; Langer and Vacanti 1993; Mattei, Giusti, and Ahluwalia 2014). The scaffold is known to be a critical determinant of cell phenotype and function and should mimic most of the properties of the native tissue extra-cellular matrix (ECM) to elicit a proper cell response (Serban and Prestwich 2008; Mattei et al. 2015). The interaction of hepatocytes with the ECM is in fact essential for the maintenance of liver specific functions (Stamatoglou and Hughes 1994), and the identification of a three-dimensional (3D) substrate which resembles the mechano-structural and physicochemical properties of the native extracellular matrix would represent a key-step towards liver tissue engineering.

Hence, it is of interest to decellularise human hepatic tissue to characterise the principal features of its ECM in order to derive quantitative design criteria to engineer biomimetic scaffolds for human hepatic cells. In this regard, we investigated the reproducibility of a given decellularisation method applied to human liver samples from 5 different donors. We started by applying our recent decellularisation method, which gave repeatable results on porcine liver samples obtained from different pigs (Mattei, Di Patria, et al. 2014), to human hepatic samples collected from 5 different donors. The decellularisation outcome was evaluated histologically using Haematoxylin-Eosin (H&E) and Alcian blue-PAS (AB/PAS) stained sections, then quantitative image analyses were performed using a purposely developed software (Magliaro et al. 2015) and the results compared to those obtained for liver samples harvested from one year old healthy pigs.

## 2 Materials and Methods

### 2.1 Liver sample collection

Normal (non-fibrotic and non-cirrhotic) human hepatic tissue was obtained from 5 patients (3 male and 2 female, mean age 68 years) undergoing liver resection for metastatic/benign liver lesions with no underlying chronic liver disease. The experiments were performed using protocols and methods in accordance with guidelines established by Italian legislation deriving from European Directives 2006/17/CE and 2006/86/CE. All patients provided their written informed consent to participate in the study after having read documentation describing the study, its motivations and their rights. The signed consent forms were registered and stored by the director of the study (Study no. 3059). The study, its methods and protocols, the consent procedure and the forms were approved by the Ethical Committee of the Azienda Ospedaliero Universita’ Pisana (approved on 21/07/2011). As indicated in the protocols, the samples received were anonymous and labelled from H1 to H5. A portion of fresh untreated tissue collected from each patient was immediately processed for histology, while the remnant part was frozen at - 20 °C within 2 hours of collection for decellularisation purposes. Frozen livers were thawed at 4 °C overnight and then shaped into 14 mm diameter – 3 mm thick cylindrical samples without Glisson’s capsule and macroscopic vasculature. Liver discs were then stored at - 20 °C until use.

Porcine hepatic tissue was harvested from 1 year-old healthy pigs (n=3) as a slaughter by-product and processed as described for human liver samples. Since porcine hepatic tissue was collected from animals killed as part of routine commercial food, no specific approval by the Local Ethical Committee was required. The results obtained for porcine liver were pooled together since animals came from the same slaughterhouse, thus no significant differences in hepatic tissue were expected or indeed observed (Marchesseau et al. 2010; Mattei, Tirella, et al. 2014; Mattei, Di Patria, et al. 2014).

### 2.2 Decellularisation procedure

Briefly, liver discs were thawed at room temperature and placed in 125 mL Nalgene square plastic bottles (Thermo Fisher Scientific, Milan, Italy). The decellularisation solution was added in a 20:1 v/w ratio with respect to the weight of liver disc samples, i.e. 40 mL solution per 2 g of liver samples (∼ 4 discs). Filled bottles were placed on an orbital shaker (SO3, Stuart Scientific, Stone, UK) at 200 rpm in a cold room at 4 °C and the decellularisation solution changed every 12 hours (day 1, PBS 1x; day 2, 1 % v/v Triton X-100 in deionised water; day 3, 12 h 0.1 % v/v Triton X-100 in deionised water + 12 h PBS 1x). All reagents were purchased from Sigma-Aldrich (Milan, Italy).

### 2.3 Histological preparation

Untreated and decellularised tissue was drop-fixed in a 10 % w/v formalin solution for 48 h. Fixed tissue was first dehydrated through graded alcohols (70, 80, 95, 100 % v/v) and embedded in paraffin. Samples were cut with a microtome (RM 2055, Leica, Heidelberg, Germany) into 5 µm sections and stained with H&E (Carlo Erba, Milan, Italy) and with Alcian Blue pH 2.5 PAS kit (Bio-Optica, Milan, Italy). In particular, n=3 different H&E and AB/PAS stained sections were examined using a Leitz Diaplan light microscope (Leitz, Wetzlar, Germany) for each of the untreated or decellularised liver sample. Each tissue section was imaged at n=5 different non-overlapping regions with a 25x objective using a digital camera (Nikon Digital Sight DS-U1, Nikon Instruments, Florence, Italy) interfaced with the NIS-Elements BR-4.13.00 software (Nikon Instruments), thus obtaining a set of n=15 images per sample/treatment/staining for subsequent quantitative histological analyses. To allow meaningful and comparative quantitative analyses both the sample preparation (e.g. slice thickness, staining procedure) and the acquisition parameters (e.g. microscope, objectives, lamp intensity, camera settings) were kept constant for all histological sections stained with the same method.

Additional 5 µm histological sections (n=3) were prepared for both untreated and decellularised porcine hepatic tissue, stained with Mallory’s trichrome and imaged as previously described for H&E and AB/PAS sections with a 10x objective in order to quantify sample collapse after the decellularisation treatment.

### 2.4 Quantitative histology

All images were analysed with HisTOOlogy, an open-source tool we developed to identify and quantify biological structures from histological sections (Magliaro et al. 2015). After loading the image to analyse and selecting the pixel size and number of dyes through a dedicated graphical user interface (GUI), the software returns objective and quantitative information on the image using a k-means clustering based dye-specific colour detection and separation algorithm. All H&E stained sections were analysed to determine the eosin covered area (ECA) and the nuclear count (NC). Briefly, the ECA was automatically quantified in the red-green-blue (RGB) colour space (see Magliaro et al. (Magliaro et al. 2015) for more details) and expressed in μm^2^ using the known image pixel size set in the GUI. The NC was evaluated by identifying the haematoxylin-segmented areas followed by object counting and expressed as number of cell nuclei per mm^2^.

AB/PAS stained sections were analysed in the cyan-magenta-yellow-black (CMYK) colour space. In particular, since AB and PAS stained regions might be co-localized (unlikely in H&E), a specific analysis of pixel intensity on both cyan and magenta channels was performed, assuming that cyan is directly related to the Alcian Blue dye and magenta to PAS. The mean pixel intensity (MPI) of cyan and magenta channels was calculated considering the total number of image pixels: variations in MPI reflect changes in the respective amount of dye.

### 2.5 Statistical analysis

Quantitative histological analyses were carried out on n=15 images per sample/treatment/staining. Differences in eosin covered area, number of cell nuclei, cyan and magenta intensity between human and porcine samples were tested using one-way ANOVA analysis followed by Tukey’s Multiple Comparison Test: untreated samples were analysed separately from the decellularised ones. For each sample, statistical differences in each of the investigated parameters before and after decellularisation were evaluated using the Student’s t-test. Statistical significance was set at *p* < 0.05.

## 3 Results

### 3.1 Decellularisation outcome

Human and pig liver disc-shaped samples were decellularised using a 3 day long immersion and agitation procedure based on non-ionic detergents, as described in Mattei et al. (Mattei, Di Patria, et al. 2014). Untreated and decellularised tissue was fixed and embedded, followed by staining with H&E and with AB/PAS to evaluate cell removal/tissue architecture and glycogen/glycosaminoglycan content, respectively. The H&E panel in Figure 1A shows a heterogeneous decellularisation outcome for human livers. Sample H5 did not appear to have been affected by the treatment, while only H4 underwent complete cell removal. Conversely, the pig livers were reproducibly (and completely) decellularised with a well-preserved tissue architecture, as previously reported (Mattei, Di Patria, et al. 2014).

**Figure 1.**
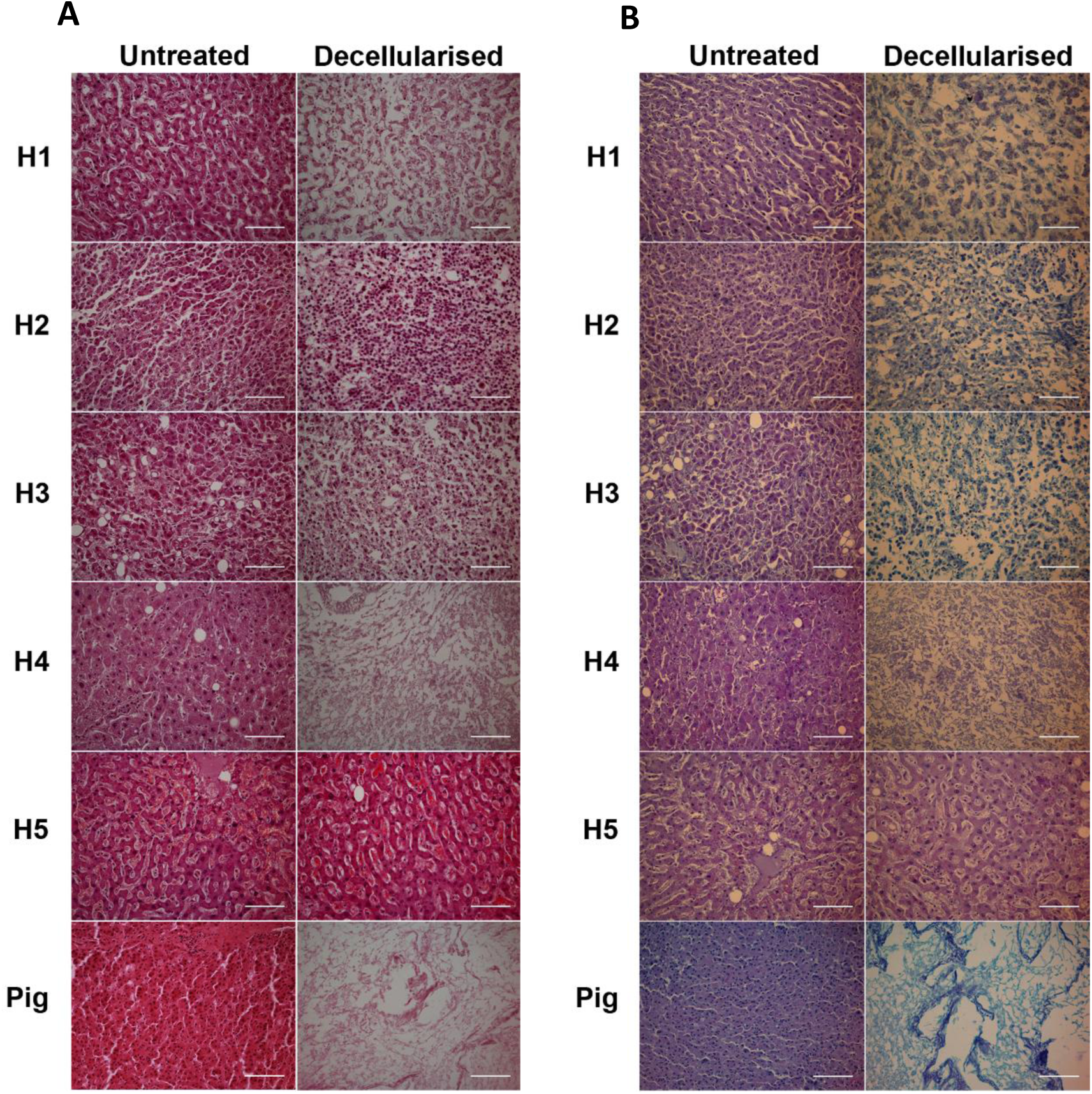
A). Haematoxylin and Eosin (H&E) stained micrographs for human (H1 to H5) and pig liver samples. Scale bar 50 µm., B). Alcian blue-PAS (AB/PAS) stained micrographs for human (H1 to H5) and pig liver samples. Scale bars 50 µm.

A decrease of the PAS-positive components (i.e. glycogen and neutral glycosaminoglycans, stained in magenta) was seen for all human samples after decellularisation, but to different extents for each donor (Fig. 1B). This decrease, which is also present in pigs, is likely due to the water-soluble nature of most of PAS-positive molecules. On the other hand, the acidic glycosaminoglycans (stained in blue) are well preserved after decellularisation, in agreement with results recently reported by Struecker et al. (Struecker et al. 2015). No visible differences in the AB/PAS stained sections were observed for sample H5, in agreement with the H&E results.

### 3.2 Sample collapse correction

A gradual collapse in hepatic lobule structure with decellularisation time was observed for pig liver samples. This collapse, which was found to be independent of the nature of the chemical detergent employed, is due to the presence of voids left by cell removal and partial ECM degradation. These regions cannot withstand mechanical loads and so result in unavoidable sample shrinking during paraffin embedding. Indeed, sample collapse due to the presence of voids which result from the decellularisation process is likely to increase the amount of a given dye or number of cell nuclei per unit area in histological sections, which is obviously meaningless. A similar artefact is encountered in many reports where quantitative values of ECM constituents (e.g., proteins, glycosaminoglycans) obtained for decellularised tissues are normalised with respect to the weight of samples after decellularisation, instead of that before decellularisation (Baptista et al. 2011; Vavken, Joshi, and Murray 2009). As a consequence, matrix components measured after decellularisation may be higher than those measured for untreated samples, making comparisons meaningless and likely privileging the more aggressive method, which removes more tissue components unselectively.

A geometric correction or “collapse” factor is proposed to account for this phenomenon and used to adjust data obtained from quantitative histological analyses that would otherwise be overestimated. The collapse factor, φ, is defined here as the ratio between the collapsed tissue area and the original un-collapsed or intact tissue area. This concept is schematised for a simple geometry in Figure 2 in which φ is equal to 0.25 (i.e. the sample collapses 4 times with respect to its original area). In particular, tissue areas were considered as rigid domains (black points in Fig. 2) that translate in space because of macroscopic sample collapse owing to the presence of voids. This assumption is valid as no significant external bulk compressive/tensile forces which may deform the tissue are exerted on the sample during histological preparation. Moreover, soft tissues are generally regarded as incompressible (Fung 1993). As a consequence, stained tissue areas in images of the decellularised samples (i.e. the 16 points in the right rectangle of Fig. 2) are generally larger than those that would be present in the absence of bulk sample collapse (i.e. the 4 points in the left rectangle of Fig. 2).

**Figure 2.**
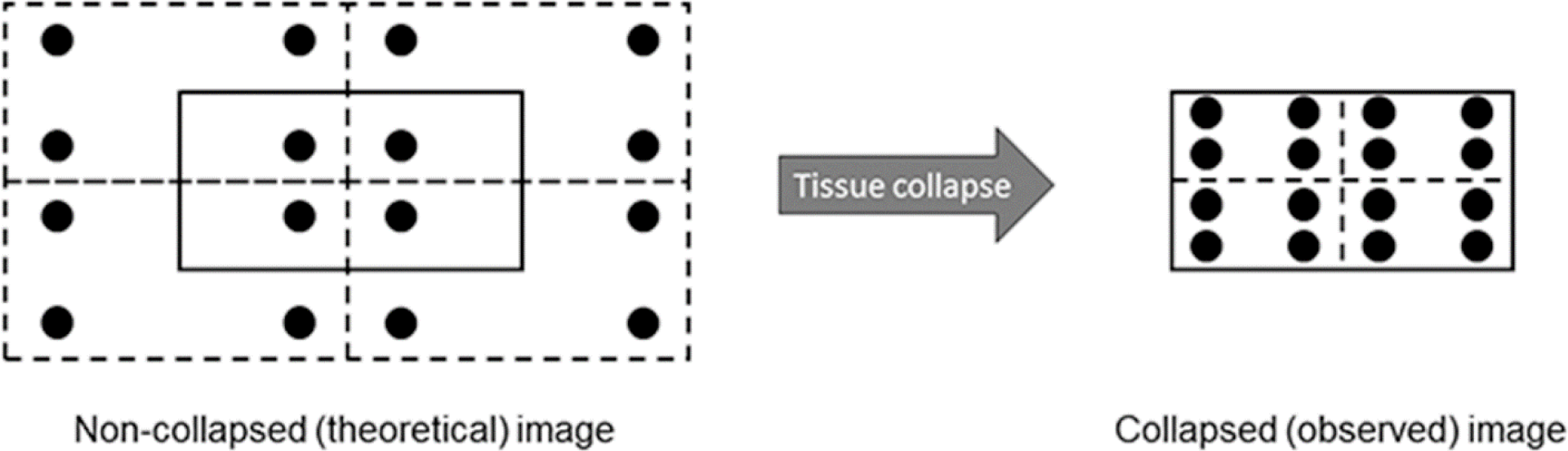
Schematic of sample collapse. In the example shown, φ = collapsed sample area / non-collapsed “original” sample area = 0.25. Thus, the sample has collapsed 4 times its original area. Specifically, tissue areas (represented by black points in figure) are regarded as rigid domains that translate in space due sample collapse. As a consequence, the collapsed sample appears to have a higher density of tissue-covered regions resulting in an overestimation of cell nuclei or ECM stain per unit area.

The collapse factor can be evaluated from histological images of untreated and decellularised tissue, by measuring characteristic areas identified using anatomical landmarks. In particular, since the boundaries of porcine liver lobules are delimited by quite thick interlobular connective tissue septa (Eurell and Frappier 2013) (clearly visible as blue lines delimiting hexagon-shaped domains in Mallory’s trichrome stained sections, Fig. 3), the average hepatic lobule area was used to quantify macroscopic collapse in decellularised samples with respect to untreated sections.

**Figure 3.**
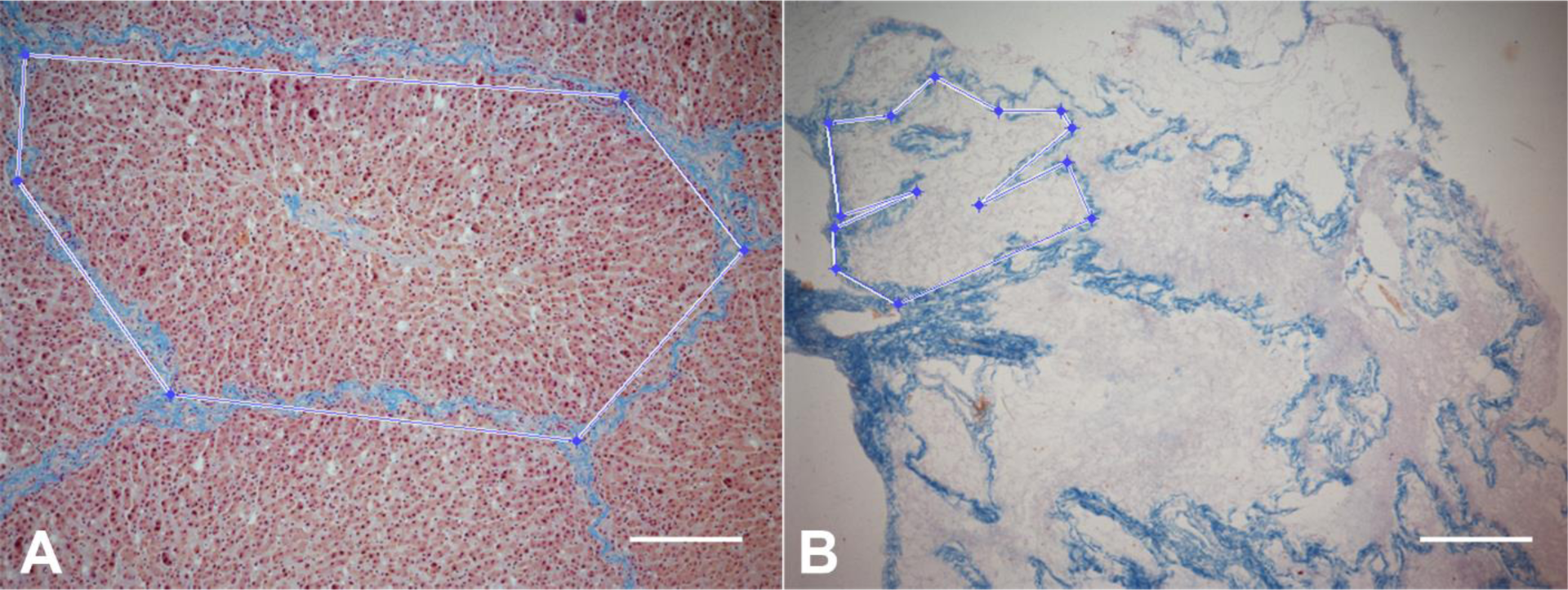
Mallory’s trichrome stained micrographs of pig liver sections. (A) Untreated and (B) decellularised samples. The deep blue colour indicates the connective tissue delimiting the hepatic lobule, the deep red in (A) stains nuclei, while the pink stain in (B) shows elastic fibres. An example of a lobule delimiting polygon is shown for both. Scale bar 100 µm.

Defining *TA* and *VA* respectively as the tissue-covered area and the void area present in a given image area (*IA*), the following “area-based” relations can be written for either the collapsed (subscript *nc*, Eq. 1) or the non-collapsed (subscript *c*, Eq. 2) case.

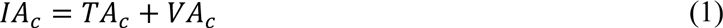

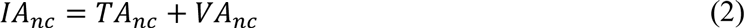

Since the optical configuration (i.e. microscope, objective, camera settings) was kept constant during image acquisition, the image area is always the same, *IA_c_* = *IA_nc_* = *IA*. As anticipated, the bulk sample collapse can be quantified through the collapse factor φ, defined as the ratio between the decellularised collapsed tissue area and the untreated un-collapsed tissue area. The collapse factor was calculated as φ = *LA_c_/LA_nc_*, where *LA_c_* and *LA_nc_* represent the average hepatic lobule area measured from histological sections of decellularised and untreated samples, respectively. Both *LA_c_* and *LA_nc_* were measured from Mallory’s trichrome stained sections using a purposely developed MATLAB script integrated in HisTOOLogy (Magliaro et al. 2015). This script allows the user to manually select the lobule contour directly on the loaded image by sequential mouse clicking and dragging, thus obtaining the polygon delimiting the lobule (Fig. 3). The area of the latter is then automatically calculated using the pixel size previously set and displayed to the user.

The quantitative histological data related to tissue-covered area *TA_c_* (e.g. amount of dye, number of cell nuclei) as well as the void area *VA_c_* for decellularized porcine samples were corrected according to Eqs. 3 and 4, thus obtaining the “true” values for decellularised samples. The corrected data can be meaningfully compared with those obtained for untreated (un-collapsed) samples. Notably, the data correction proposed here can be used either in case of bulk sample collapse (where 0 < φ < 1) or swelling (φ > 1).

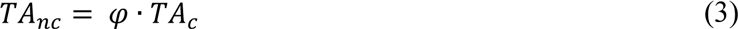

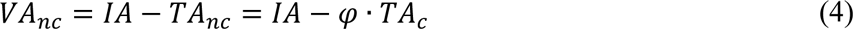

The collapse factor between decellularised and untreated porcine hepatic tissue was φ = 0.56 ± 0.08, and was used to correct quantitative histological data in tissue-covered areas (as outlined in Methods). Unlike porcine hepatic tissue, neither the connective tissue septa outlining hepatic lobules nor other visible anatomical landmarks are present in human liver (Matthews and Martin 1971), making it difficult to determine lobule areas and calculate φ as previously described. However, this is not a limitation for the present study, since no macroscopic sample collapse was observed between treated and untreated human liver sections by expert biologists, likely due to the poor effect of the decellularisation treatment of all but one sample of human hepatic tissue.

### 3.3 Quantitative image analysis

The results obtained from quantitative histological analyses of either untreated or decellularised human and porcine samples are reported in Figure 4.

**Figure 4.**
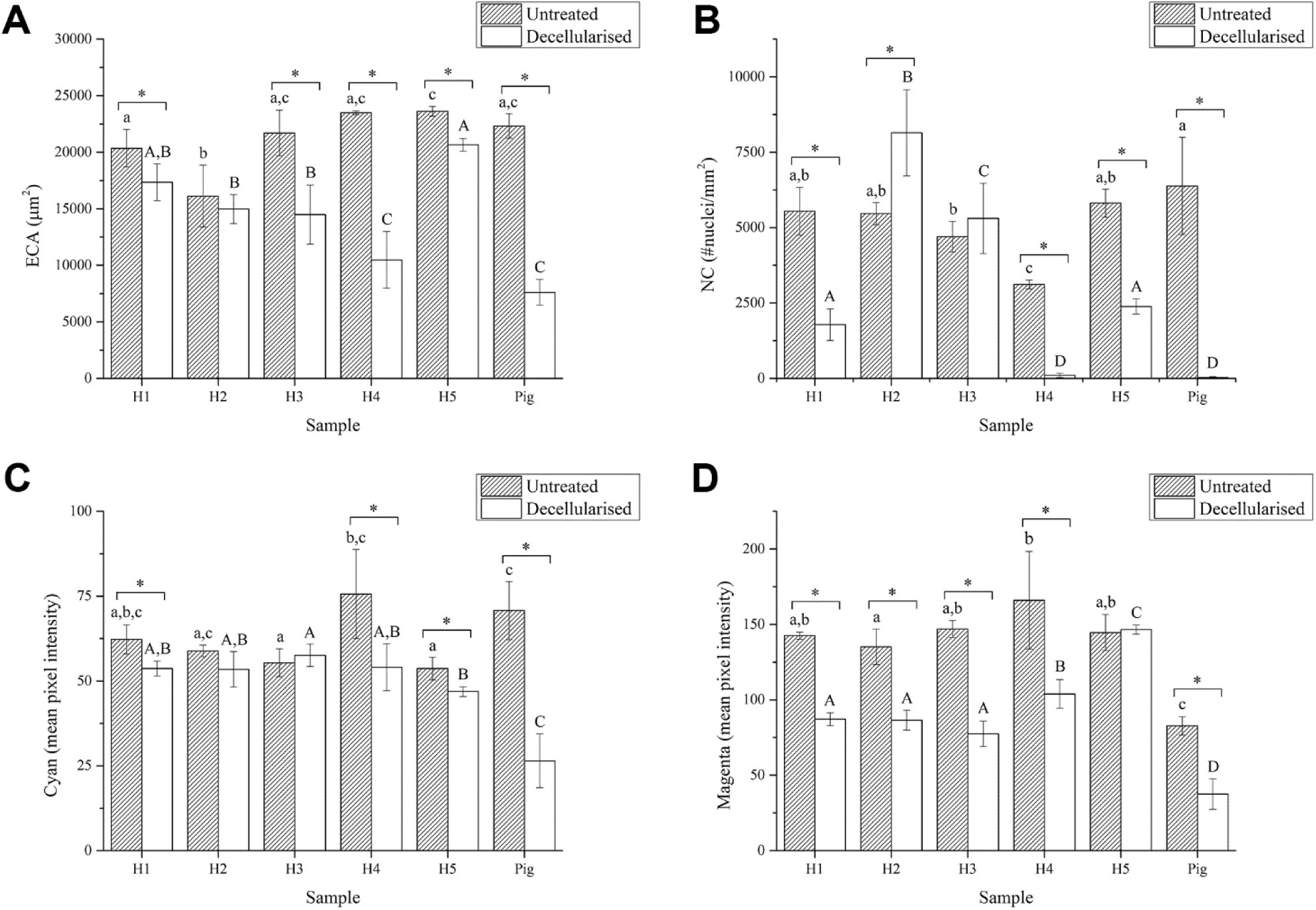
Quantitative histological analyses. (A) Eosin covered area, (B) Number of cell nuclei, (C) Cyan mean pixel intensity, (D) Magenta mean pixel intensity. Different letters indicate significant differences between groups (one-way ANOVA, p < 0.05): lowercase letters are referred to ANOVA results of untreated samples, while capital letters denote ANOVA results of decellularised samples. All pig liver data are corrected for the collapse factor. Statistical differences in each of the investigated parameter prior and after the decellularisation treatment were evaluated for each sample (Student’s t-test, p < 0.05) and denoted by an asterisk.

In particular, the eosin covered area (ECA) was found to be similar in most of the untreated human and porcine samples (Fig. 4A). Overall, the decellularisation treatment resulted in a significant decrease of the ECA (except for sample H2), as expected since this parameter is directly related to the sample total protein content (Magliaro et al. 2015) which generally decreases after decellularisation (Mattei, Di Patria, et al. 2014). However, the variability in ECA values obtained for human samples after decellularisation suggest that the outcome of the decellularisation process is donor-dependent.

The number of cell nuclei per mm^2^ (NC) decreases after decellularisation, as expected (Fig. 4B). An increase in the NC was observed for the sample H2, probably because of some sample collapse after decellularisation. Although the decrease in NC was significant for H1, H4, H5 and pig samples, complete cell removal was achieved only in case of H4 and pig, with more than 96.5 % and 99.5 % of cell nuclei removed, respectively. Again, these results confirm that the decellularisation outcome is donor-dependent.

The cyan mean pixel intensity (MPI) was found to be similar in the untreated human and pig samples (Fig. 4B). As observed for the ECA and the NC, the decellularisation treatment resulted in a decrease in cyan MPI, which was significant for H1 (13.7 %), H4 (9.1 %) and H5 (28.5 %) human samples as well as for pig (62.6 %). Assuming that cyan MPI is related to Alcian Blue dye, these results suggest that acidic glycosaminoglycans are fairly preserved after decellularisation (> 70 % for humans and > 35 % for pig), in agreement with visual inspection of Figure 1B.

Untreated human samples exhibited similar magenta MPIs, significantly higher than that observed for pig (Fig. 4D). The magenta MPI decreases significantly after decellularisation for both human (∼ 40 % on average, but with considerable scatter, except sample H5) and pig samples (54.7 %). As the magenta MPI is related to PAS, these results indicate that the decellularisation treatment causes a decrease of PAS-positive components (i.e. glycogen and neutral glycosaminoglycans), which is generally higher than that observed for the less water soluble acidic glycosaminoglycans.

## 4 Discussion

Donor livers suitable for transplant are chronically in short supply, and will become even scarcer as life expectancy increases. To meet the demand, tissue engineered livers could be developed using novel biomaterials which recapitulate the microarchitecture and cell adhesive environment of native liver ECM. In fact, the extracellular matrix is thought to be the ideal scaffold for tissue engineering. Starting from healthy organs judged unsuitable for transplant because of prolonged graft cold ischaemic time, the presence of extra-hepatic malignancy or other important extra-hepatic comorbidities in donors or recipients, whole human liver decellularisation for the generation of hepatic scaffolds has been reported (Mazza et al. 2015). However, as the goal of the study was to obtain bio-artificial liver tissue by employing decellularised human ECM liver scaffolds, the authors did not quantify the features of the scaffolds nor analyse inter-donor differences and pooled data from 3 different human livers together. Moreover, although their results are of scientific relevance, the availability of healthy human liver unsuitable for transplant is destined to diminish thanks to new techniques for prolonging organ preservation after death, such as extracorporeal membrane oxygenation (ECMO) support (Balsorano et al. 2015; Carter et al. 2014; Rojas-Peña et al. 2014). Therefore, methods that do not rely on the use of healthy human tissue are needed to overcome the shortage of donor organs.

As decellularised human liver is considered an exemplar for designing the ideal substrate for hepatic regeneration, we sought to identify a unique set of structural and biochemical characteristics of the human liver ECM in order to design a generic biomimetic hepatic scaffold for humans. To this end, we applied our recent decellularisation method, which gave repeatable results on porcine liver samples obtained from different pigs (Mattei, Di Patria, et al. 2014), to human hepatic samples collected from 5 different donors. The outcomes in terms of cell removal, protein content, and glycosaminoglycan content per unit area were compared between donors and resulted in high donor-to-donor variability. We also applied a more aggressive ionic decellularisation method replacing 1% Triton X-100 with 0.1% SDS (i.e. I3 protocol described in Mattei et al. (Mattei, Di Patria, et al. 2014)) to all samples. Although this method gave reproducible results with pigs, once again the outcomes for human hepatic tissue were still highly variable, as shown in Fig 5.

**Figure 5.**
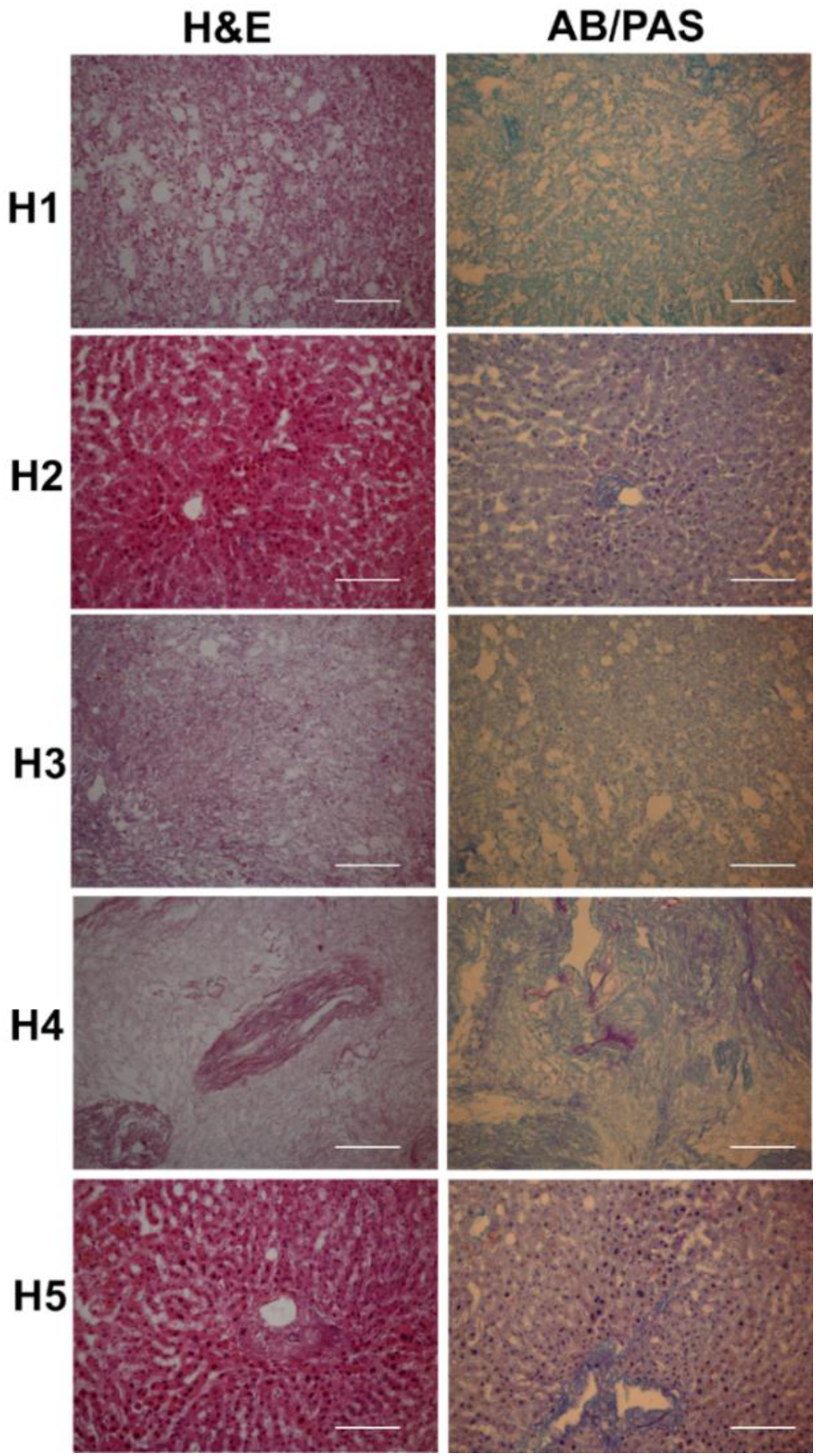
Haematoxylin and Eosin (H&E) and Alcian blue-PAS (AB/PAS) stained micrographs for human liver samples H1 to H5, decellularized using SDS. The decellularization protocol was: day 1, PBS 1x; day 2, 0.1 % v/v SDS in deionised water; day 3, 12 h 0.1 % v/v Triton X-100 in deionised water followed by 12 h in PBS 1x. Scale bar 50 µm.

In conclusion, given the high level of heterogeneity between donors, human liver ECM from different donors cannot be reproducibly decellularised using a single cell-removal protocol. Of course the decellularisation protocol could be adapted for each liver so as to achieve 100% cell removal, but if the aim is to derive ideal donor-independent design criteria for engineering a generic “one-size fits all” human hepatic scaffold, the adaptation cannot be donor-dependent. One could also argue that the protocols investigated here are not suitable for our samples, and a more aggressive protocol is necessary for complete cell removal in human livers. However, the more aggressive the decellularisation treatment, the less faithful or “biomimetic” is the matrix to the native ECM microenvironment (Peloso et al. 2015). Thus, decellularised human liver cannot be used as a template to derive quantitative design criteria for engineering a generic hepato-mimetic scaffold. Our results suggest that scaffold design criteria should be tailored to the recipient, in agreement with emerging personalised medicine strategies (Schleidgen et al. 2013). Alternatively, porcine liver dECM, whose mechano-structural and physicochemical features have been shown to be donor-independent (Marchesseau et al. 2010; Mattei, Tirella, et al. 2014; Mattei, Di Patria, et al. 2014), may represent an attractive candidate as a template for deriving scaffold specifications, since human primary hepatocytes exhibit good metabolic competency when cultured on porcine liver-derived ECM scaffolds, either in the form of 3D porous matrices (Lang et al. 2011) or hydrogels (Sellaro et al. 2010). Thus, an in-depth quantitative mechano-structural analysis of porcine-derived hepatic dECM could be performed to establish a unique set of design specifications for engineering scaffolds for human livers.

## 6 Author Contributions

GM, CM and AA designed the research; GM, CM and AP performed the research; GM, CM, AP and AA analyzed the data; GM and AA wrote the paper. All authors read and approved the final manuscript.

## 7 Acknowledgments

The work leading to these results has received funding from the European Union Seventh Framework Programme (FP7/2007-2013) under grant agreement 304961 (ReLiver).

